# Unsupervised integration of single-cell multi-omics datasets with disparities in cell-type representation

**DOI:** 10.1101/2021.11.09.467903

**Authors:** Pinar Demetci, Rebecca Santorella, Björn Sandstede, Ritambhara Singh

## Abstract

Integrated analysis of multi-omics data allows the study of how different molecular views in the genome interact to regulate cellular processes; however, with a few exceptions, applying multiple sequencing assays on the same single cell is not possible. While recent unsupervised algorithms align single-cell multi-omic datasets, these methods have been primarily benchmarked on co-assay experiments rather than the more common single-cell experiments taken from separately sampled cell populations. Therefore, most existing methods perform subpar alignments on such datasets. Here, we improve our previous work Single Cell alignment using Optimal Transport (SCOT) by using unbalanced optimal transport to handle disproportionate cell-type representation and differing sample sizes across single-cell measurements. We show that our proposed method, SCOTv2, consistently yields quality alignments on five real-world single-cell datasets with varying cell-type proportions and is computationally tractable. Additionally, we extend SCOTv2 to integrate multiple (*M* ≥ 2) single-cell measurements and present a self-tuning heuristic process to select hyperparameters in the absence of any orthogonal correspondence information.

**Available at:** http://rsinghlab.github.io/SCOT.

## 1 Introduction

The ability to measure multiple aspects of the single-cell offers the opportunity to gain critical biological insights about cell development and diseases. However, many existing single-cell sequencing technologies cannot be simultaneously applied to the same cell, resulting in multi-omics datasets sampled from distinct cell populations. While these measurements can be analyzed separately, integrating them prior to analysis can help explain how different molecular views interact and regulate cellular functions. Unfortunately, single-cell assays that measure different molecular aspects in separately sampled cell populations lack direct sample–sample and feature–feature correspondences across these measurements. This lack of correspondences makes it hard to use integration methods that require some shared information to perform single-cell alignment [1]. Therefore, *unsupervised* single-cell multi-omics data alignment methods are crucial for integrative single-cell data analysis.

Several unsupervised methods [1–4], including our previous work, SCOT [5], have shown state-of-the-art performance for integrating different single-cell measurement domains. Since these methods were mainly evaluated on real-world co-assay datasets (with 1–1 correspondence between cells across domains), our understanding of their performance on datasets obtained from experiments that are not co-assays is limited. Such experiments perform separate sampling to measure distinct genomic features, like gene expression and 3D chromatin conformation. Therefore, their datasets can consist of varying proportions of cell-types across different measurements, creating cell-type imbalance and lacking 1–1 cell correspondences. We hypothesize that alignment methods that perform well on co-assay datasets may not effectively handle the differences in cell-type proportions of the commonly available non-co-assay datasets. Indeed, a recent method, Pamona [6], extended our SCOT framework and used partial Gromov-Wasserstein (GW) optimal transport to allow for missing or underrepresented cell-types in one domain when performing alignment. The paper showed that current integration methods [1, 2, 4, 5] tend to perform worse under such settings.

We present SCOTv2, a novel extension of SCOT that can effectively align both co-assay and non-co-assay datasets using a single framework. It uses *unbalanced* GW optimal transport to align datasets with disproportionate cell-types while only introducing one additional hyperparameter. This unbalanced framework relaxes the constraint that each point must be mapped with its original mass during the optimal transport. Specifically, an underrepresented cell-type in one domain can be transported with more mass to match the proportion of that cell-type in the other domain and vice-versa. The SCOTv2 framework is summarized in Figure 1. We demonstrate that SCOTv2 aligns datasets with imbalance in cell-type representations better than state-of-the-art baselines and computationally scales as well as the fastest methods. Furthermore, we extend SCOTv2 to integrate single-cell datasets with more than two measurements, making it a multi-omics alignment tool. We perform alignments of five real-world single-cell datasets, with both simulated and natural cell-type imbalance as well as two and more than two domains (*M* ≥ 2), demonstrating SCOTv2’s applicability across a wide range of scenarios. Finally, similar to the previous version, we present a self-tuning heuristic process to select hyperparameters for SCOTv2 without any corresponding information like cell-type annotations or matching cells or features in truly unsupervised settings.

**Figure 1:**
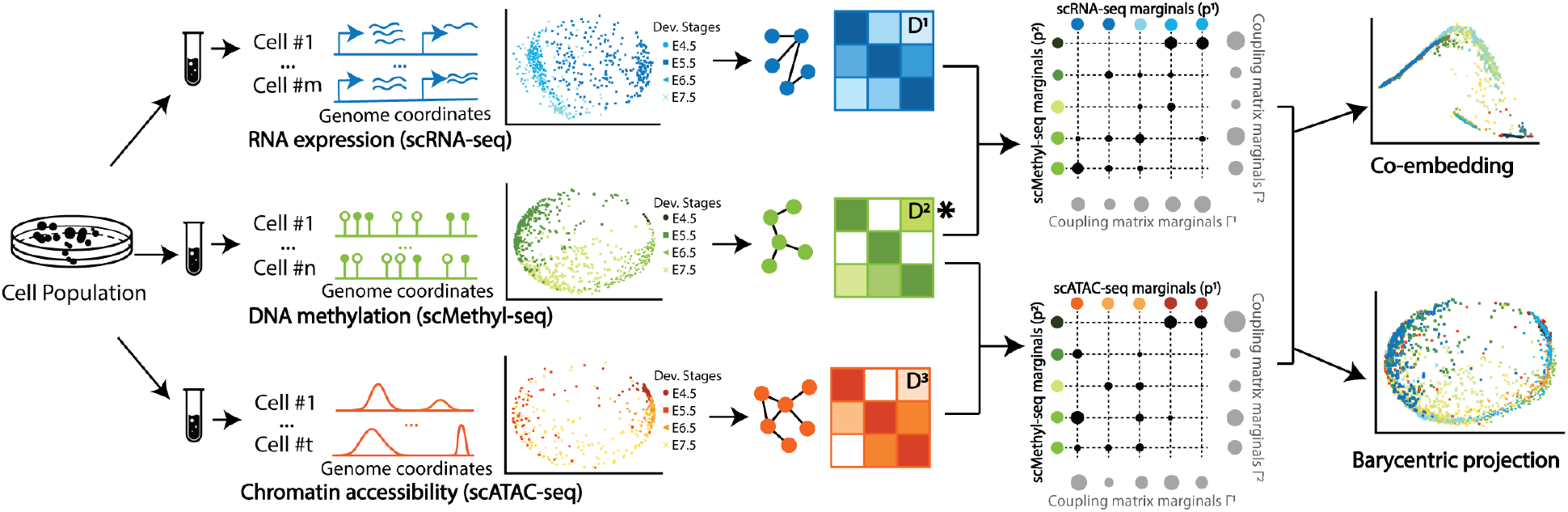
Overview of SCOTv2 on scNMT-seq dataset [7], which contains unbalanced cell-type representation across three domains - RNA expression, chromatin accessibility, and DNA methylation. SCOTv2 selects an anchor domain (denoted with **\***) and aligns other measurements to it. First, it computes intra-domain distances matrices *D*^*m*^ for *m* = 1, 2, 3, which are used to solve for correspondence matrices between the anchor and other domains. The circle sizes in the matrices depict the magnitude of the correspondence probabilities or how much mass to transport. Unbalanced GW relaxes the mass conservation constraint, so the transport map does not need to move each point with its original mass. Finally, it either co-embeds the domains into a common space or uses barycentric projections to project them onto the anchor domain.

## 2 Method

Optimal transport finds the most cost-effective way to move data points from one domain to another. One can imagine it as the problem of moving a pile of sand to fill in a hole through the least amount of work. Our previous framework SCOT [5] uses Gromov-Wasserstein optimal transport, which preserves local geometry when moving data points from one domain to another. The output of SCOT is a matrix of probabilities that represent how likely it is that data points from one modality correspond to data points in the other.

Here, we reintroduce the SCOT formulation to integrate *M* domains (or single-cell measurements) 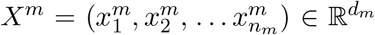 for *m* = 1, … *M* with *n*_*m*_ data points (or cells) each. For each dataset, we define a marginal distribution *p*^*m*^, which can be written as an empirical distribution over the data points:

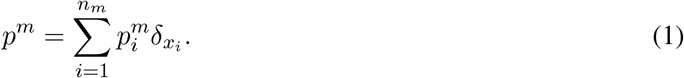

Here, 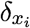 is the Dirac measure. For SCOT, we choose these distributions to be uniform over the data.

Gromov-Wasserstein optimal transport performs the transport operation by comparing distances between samples rather than directly comparing the samples themselves [8]. Therefore, for each dataset, we compute the intra-domain distance matrix *D*^*m*^. Next, we construct *k*-NN graphs based on correlations between data points and use Dijkstra’s algorithm to compute the shortest path distance on the graph between each pair of nodes. Finally, we connect all unconnected nodes by the maximum finite distance in the graph and set *D*^*m*^ to be the matrix resulting from normalizing the distances by this maximum.

For two datasets and a given cost function *L* : ℝ × ℝ → ℝ, we compute the fourth-order tensor 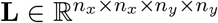, where 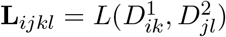. Intuitively, *L* quantifies how transporting a pair of points 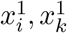 onto another pair across domains, 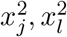, distorts the original intra-domain distances and helps to preserve local geometry. Then, the discrete Gromov-Wasserstein problem between *p*^1^ and *p*^2^ is,

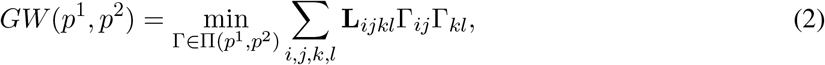

where Γ is a coupling matrix from the set:

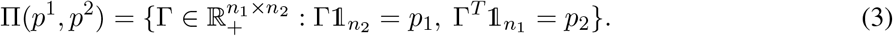

One of the advantages of using optimal transport is the probabilistic interpretation of the resulting coupling matrix Γ, where the entries of the normalized row 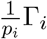 are the probabilities that the fixed data point *x*_*i*_ corresponds to each *y*_*j*_. Each entry Γ_*ij*_ describes how much of the mass of *x*_*i*_ should be mapped to *y*_*j*_.

To make this problem more computationally tractable, we solve the entropically regularized version:

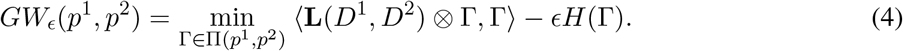

where *ϵ* > 0 and *H*(Γ) is the Shannon entropy defined as 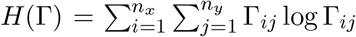. Larger values of *ϵ* make the problem more convex but also lead to a denser coupling matrix, meaning there are more correspondences between samples. In SCOT, we use the cost function *L* = *L*_2_.

### 2.1 Unbalanced Optimal Transport of SCOTv2

Our proposed solution to align datasets with different numbers of samples or proportions of cell-types is to use unbalanced optimal transport, which adds divergence terms to allow for mass variations in the marginals [9, 10]. We follow Séjourné *et al* [10], and use the Kullback-Leibler divergence,

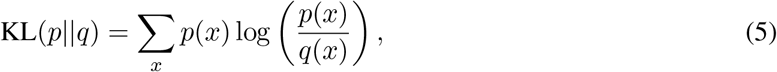

to measure the difference between the marginals of the coupling Γ and the input marginals *p*^1^ and *p*^2^. Thus, we solve the unbalanced GW problem:

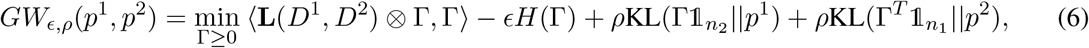

where *ρ* > 0 is a hyperparameter that controls the marginal relaxation. When *ρ* is large, the marginals of Γ should be close to *p*^1^ and *p*^2^, and when *ρ* is small, the marginals of Γ may differ more, allowing each point to transport with more or less mass than it originally had. See Supplementary Algorithm 2 for details.

### 2.2 Extending SCOTv2 for Multi-Domain Alignment

To align more than two datasets (*M* > 2), we use one domain as an anchor to align the other domains. The anchor should be the domain with the clearest biological structures, for example, a dataset with the best-defined cell-type clusters. We propose selecting the anchor via the kNN graph used to compute *D*^*m*^. For every node 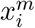 in the graph, we calculate the average of the *k* neighboring node values 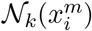. Next, we measure the difference between this average and the true value of the node. This difference reflects how well the averaged neighborhood represents the given node. We then average these differences across the graph and select the domain with the lowest averaged difference as the anchor. Intuitively, we select the anchor whose kNN graph best reflects its dataset. Suppose *X*^1^ is the anchor dataset. Then, for *m* = 2, 3, …, *N*, we compute the coupling matrix Γ^*m*^ according to Equation 4.

To have all of the datasets aligned in the same domain, we can either use barycentric projection to project each *X*^*m*^ for *m* = 2, 3, …, *M* onto *X*^1^ or find a shared embedding space as described in Section 2.3. In the first iteration of SCOT, we used a barycentric projection to align and project one dataset onto the other. Due to the marginal relaxation, we now search for a non-negative *n*_1_ × *n*_*m*_ dimensional matrix Γ instead of Γ ∈ Π(*p*^1^, *p*^*m*^). Because of this change, the adjusted barycentric projection is:

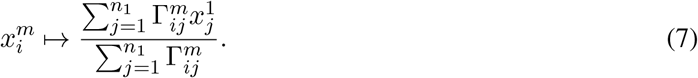

### 2.3 Embedding with the Coupling Matrix

Other methods such as MMD-MA and UnionCom align datasets by embedding them into a common latent space of dimension *p* ≤ min_*m*=1,…,*M*_ *d*_*m*_. Here *d*_*m*_ represents the original dimension size of measurement (or domain) *m*. Embedding the datasets in a new space often leads to a better alignment as it introduces the additional benefits of dimension reduction, allowing more meaningful structures in the datasets such as cell-types to be more prevalent. Due to these benefits, we also enable the embedding option through a modification of the t-SNE method proposed by UnionCom [1]. For each domain *m*, we compute *P*^*m*^, an *n*_*m*_ × *n*_*m*_ cell-to-cell transition matrix; each entry 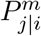 is the conditional probability that a data point 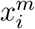 would pick 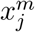 as its neighbor when chosen according a Gaussian distribution centered at 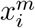. Similarly, for the lower-dimensional embeddings, we compute a cell-to-cell probability matrix *Q*^*m*′^ through a Student-t distribution. The full descriptions of *P*^*m*^ and *Q*^*m*′^ are given in Supplementary Section S1.

Then, to jointly embed all domains through the anchor domain *X*^1^, the optimization problem is:

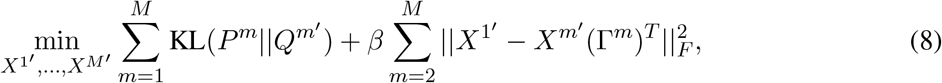

where *X*^*m*′^ is the lower dimensional embedding of *X*^*m*^, and Γ^*m*^ is the coupling matrix from solving Equation 6 for *m* = 2, …, *M*. These two terms seek to find an embedding that both preserves the local geometry in the original domain and aligns the domains according to the correspondence found by GW. The intuition behind the term KL(*P*^*m*^||*Q*^*m*′^) is very similar to that of GW; if two points have a high transition probability in the original space, then they should also have a high transition probability in the latent space. The term 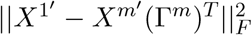 measures how well aligned the new embeddings *X*^1′^ and *X*^*m*′^ are according to the prescribed coupling matrix Γ^*m*^. Finally, *β* > 0 controls the trade-off between preserving the original geometry with the KL term and enforcing the alignment found with GW. We solve this optimization problem using gradient descent from UnionCom with a default latent space dimension size *p* = 3 [1]. The overall SCOTv2 method is presented as Supplementary Algorithm 3.

### 2.4 Heuristic process for self-tuning hyperparameters

SCOTv2 has three hyperparameters: (1) *k* for the number of neighbors to consider in nearest neighbor graphs, (2) the weight of the entropic regularization term, *ϵ*, and (3) the coefficient of the mass relaxation constraint, *ρ*. The barycentric projection of one domain onto another does not require any hyperparameters. However, jointly embedding the domains in a latent space requires selecting the dimension *p*.

Ideally, orthogonal correspondence information such as 1–1 correspondences and cell-type labels can guide hyperparameter tuning as validation. However, such information is hard to obtain in most cases. First, no validation data on cell-to-cell correspondences exists for non-co-assay datasets. Second, it is challenging to infer cell-types for certain sequencing domains such as 3D chromatin conformation. Lastly, the cell-type annotations may not always agree across single-cell domains.

We provide a heuristic to self-tune hyperparameters in the completely unsupervised setting. We first choose a *k* for the neighborhood graphs that yields a high average correlation value between the neighborhood predicted values and measured genomic values of the graph nodes. This step is the same as the one used to select the anchor domain for multi-omics alignment in Section 2.2. Next, we choose *ϵ* and *ρ* values that minimize the Gromov-Wasserstein distance between the aligned datasets. Algorithm 1 gives the details of this procedure.

#### Algorithm 1: Unsupervised hyperparameter search procedure

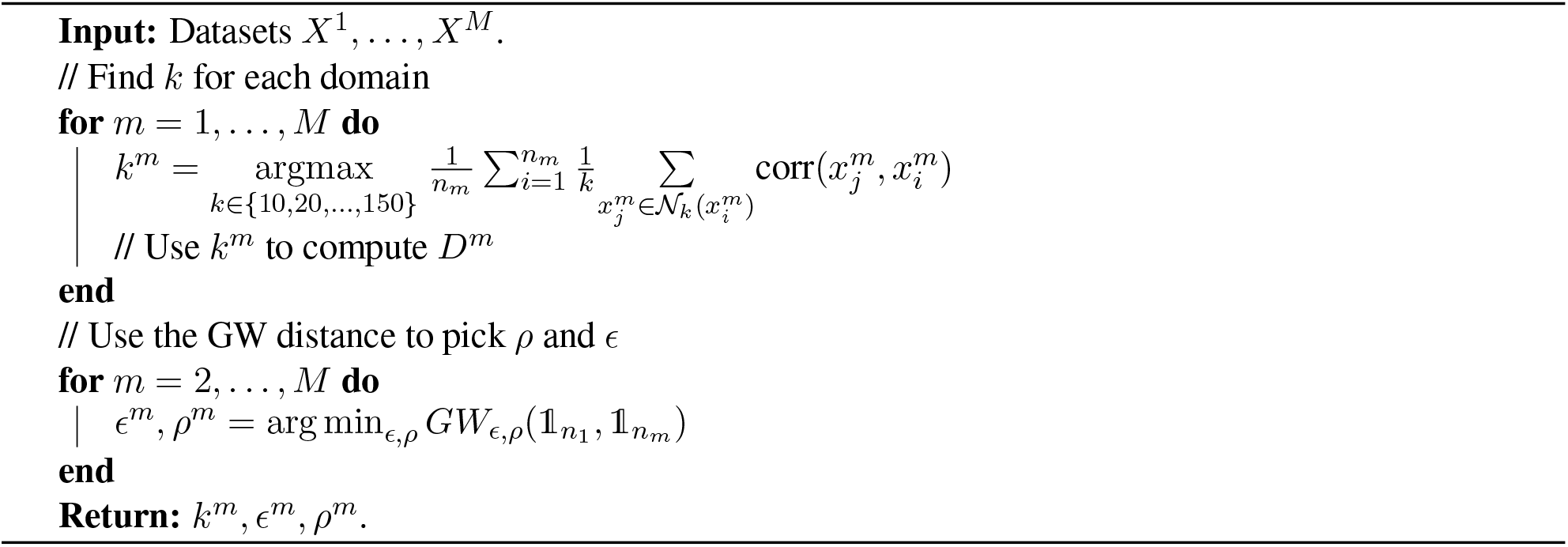

## 3 Experimental Setup

### 3.1 Datasets

We evaluate SCOTv2 on single-cell datasets with disproportionate cell-types using two schemes. (1) We subsample different cell-types in co-assay datasets to simulate cell-type representation disparities between sequencing modalities. (2) We select real-world separately sequenced single-cell multi-omics datasets, which lack 1–1 cell correspondences and have different cell-type proportions across modalities due to the sampling procedure. Additionally, we present results on the original co-assay datasets with 1–1 cell correspondence to demonstrate the flexibility of SCOTv2 across balanced and unbalanced single-cell datasets.

#### 3.1.1 Co-assay single-cell datasets with 1–1 cell correspondence

We use three co-assay datasets to validate our model, sequenced by SNARE-seq, scGEM, and scNMT technologies. SNARE-seq is a two-modality sequencing technology that simultaneously captures the chromatin accessibility and transcriptional profiles of cells [11]. This dataset contains a total of 1047 cells from four cell lines: BJ (human fibroblast cells), H1 (human embryonic cells), K562 (human erythroleukemia cells), and GM12878 (human lymphoblastoid cells) (Gene Expression Omnibus access code: GSE126074). We follow the same data preprocessing steps outlined by Chen *et al*. [11]. The scGEM technology is a three-modality sequencing technology that profiles the genetic sequence, gene expression, and DNA methylation states in the same cell [12]. The dataset we use is derived from human somatic cell samples undergoing conversion to induced pluripotent stem cells (Sequence Read Archive accession code SRP077853) [12]. We access the preprocessed data provided by Welch *et al*. [13], which only contains the gene expression and DNA methylation modalities ^1^. The dataset sequenced by scNMT-seq method [14] contains three modalities of genomic data: gene expression, DNA methylation, and chromatin accessibility, from mouse gastrulation samples, going through the Carnegie stages of vertebrate development (Gene Expression Omnibus access code: GSE109262). We access the preprocessed data through the scripts^2^ provided by the authors. While the SNARE-seq and scGEM datasets contain the same number of cells across measurements, scNMT-seq modalities contain different cell-type proportions after preprocessing due to varying noise levels in measurements. Supplementary Table S1 lists the number of cells belonging to different cell-types in each domain for scNMT-seq dataset.

#### 3.1.2 Single-cell datasets with simulated cell-type imbalance

To test alignment performance sensitivity to different levels and types of cell-type proportion disparities across modalities, we generate simulation datasets by subsampling SNARE-seq and scGEM co-sequencing datasets in two ways. (1) We remove a cell-type from one modality. (2) We reduce the proportion of a celltype in one modality by subsampling it at 50% and another cell-type in the other modality by subsampling it at 75%. We simulate this setting to test how the alignment methods will behave when multiple cell-types have disproportionate representation at different levels (for example, half or quarter percentage of cell-types missing) across modalities.

For these cases, we uniformly pick at random which cell-type to subsample or remove. Specifically, for scGEM in simulation case (1), we remove “d16T+” cells in the DNA methylation domain while retaining the original gene expression domain, and remove the “d24T+” cells in the gene expression domain while retaining the original DNA methylation domain. For the SNARE-seq dataset, we remove “GM” cells in the gene expression domain and “K562” in the chromatin accessibility domain. In simulation case (2), we subsample the “d8” cluster of the scGEM dataset at 75% in the gene expression modality and the “d16T+” cluster at 50% in the DNA methylation modality. For SNARE-seq, we subsample the “H1” cluster at 75% and the “K562” cluster at 50% in the gene expression and chromatin accessibility domains, respectively.

#### 3.1.3 Single-cell datasets without 1—1 correspondences

We also align non-co-assay datasets, containing separately sequenced single-cell -omic measurements. Bonora *et al*. generated the first dataset we use, “sciOmics” [15]. This dataset consists of sciRNA-seq, sciATAC-seq, and sciHiC measurements, capturing gene expression, chromatin accessibility, and 3D chromosomal conformation profiles of mouse embryonic stem cells undergoing differentiation. The measurements were taken at five stages: days 0, 3, 7, 11, and as fully differentiated neural progenitor cells (NPCs). The second non-co-assay dataset, “MEC,” contains gene expression and chromatin accessibility measurements taken using the 10X Chromium scRNA-seq and scATAC-seq technologies on mouse mammary epithelial cells (MEC). Since each modality consists of separately sampled cell populations, these contain disparate cell-type proportions across modalities. Supplementary Table S1 lists the number of cells belonging to different cell-types in each domain for sciOmics and MEC datasets.

### 3.2 Evaluation metrics and baseline methods

Although most of the datasets lack 1–1 cell correspondences, we can evaluate alignment using cell-type labels through label transfer accuracy (LTA) as in [1, 5, 6]. This metric assesses the clustering of cell-types after alignment by training a *k*NN classifier on a training set (50% of the aligned data) and then evaluates its predictive accuracy on a test dataset (the other 50% of the aligned data). Higher values correspond to better alignments, indicating that cells that belong to the same cell-type are aligned close together after integration. We benchmark our method against the current unsupervised single-cell multi-omic alignment methods, Pamona [6], UnionCom [1], MMD-MA [16], bindSC [3], Seuratv4 [4], and the previous version of SCOT, which performs alignment without the KL term [5]. Pamona [6], as previously discussed, uses partial Gromov-Wasserstein (GW) optimal transport to align single-cell datasets. UnionCom [1] performs unsupervised topological alignment through a two-step procedure that first finds a correspondence between the domains, considering both global and local geometries with a hyperparameter to control the trade-off between them, and then embeds them in a new shared space. MMD-MA [16] uses the maximum mean discrepancy (MMD) measure to align and embed two datasets in a new space. BindSC [3] requires the users to bring input datasets to the gene expression feature space by constructing a gene activity score matrix for the epigenomic domains, then finds a correspondence matrix between samples through bi-order canonical correspondence analysis (bi-CCA), and jointly embeds the domains into a new space. Finally, Seuratv4 [4] also requires gene activity score matrices for epigenomic domains and then identifies correspondence anchors via CCA. Based on these anchors, it imputes one genomic domain based from the other domain and co-embeds them into a shared space using UMAP.

Since bindSC and Seurat v4 require the creation of gene activity score matrices for epigenomic datasets, they might be more difficult to use with certain sequencing domains. For instance, gene activity scoring is challenging for 3D chromosomal conformation. Of all the selected baselines, only Pamona and UnionCom can align more than two domains, so we only use them as baselines for experiments with multiple domains (*M* > 2). For each benchmark, we define a hyperparameter grid of similar granularity and perform extensive tuning (see Supplementary Section S4). We report the alignment results with the best performing hyperparameter combinations in Section 4.1.

## 4 Results

### 4.1 SCOTv2 gives high-quality alignments consistently across all single-datasets

We first present the alignment results for real-world co-assay datasets with simulated cell-type imbalance. Figure 2 (A) visualizes the barycentric projection alignments performed by SCOTv2 plotted as 2D PCA for SNARE-seq and scGEM datasets, respectively. We use barycentric projection for visualization purposes for the ease of comparison with the original domains, plotted in Supplementary Figure S1. Here, we integrate datasets under three different settings described in the previous section: (1) Balanced datasets (or “full datasets” with no subsampling), (2) Missing cell-type in the epigenomic domains, and (3) Subsampled cells in both domains (one cell-type at 50% in the epigenomic domains and another cell-type at 75% in the gene expression domains). We include alignment results on the full datasets with 1–1 sample correspondences to ensure that SCOTv2 performs well for balanced cases as well.

**Figure 2:**
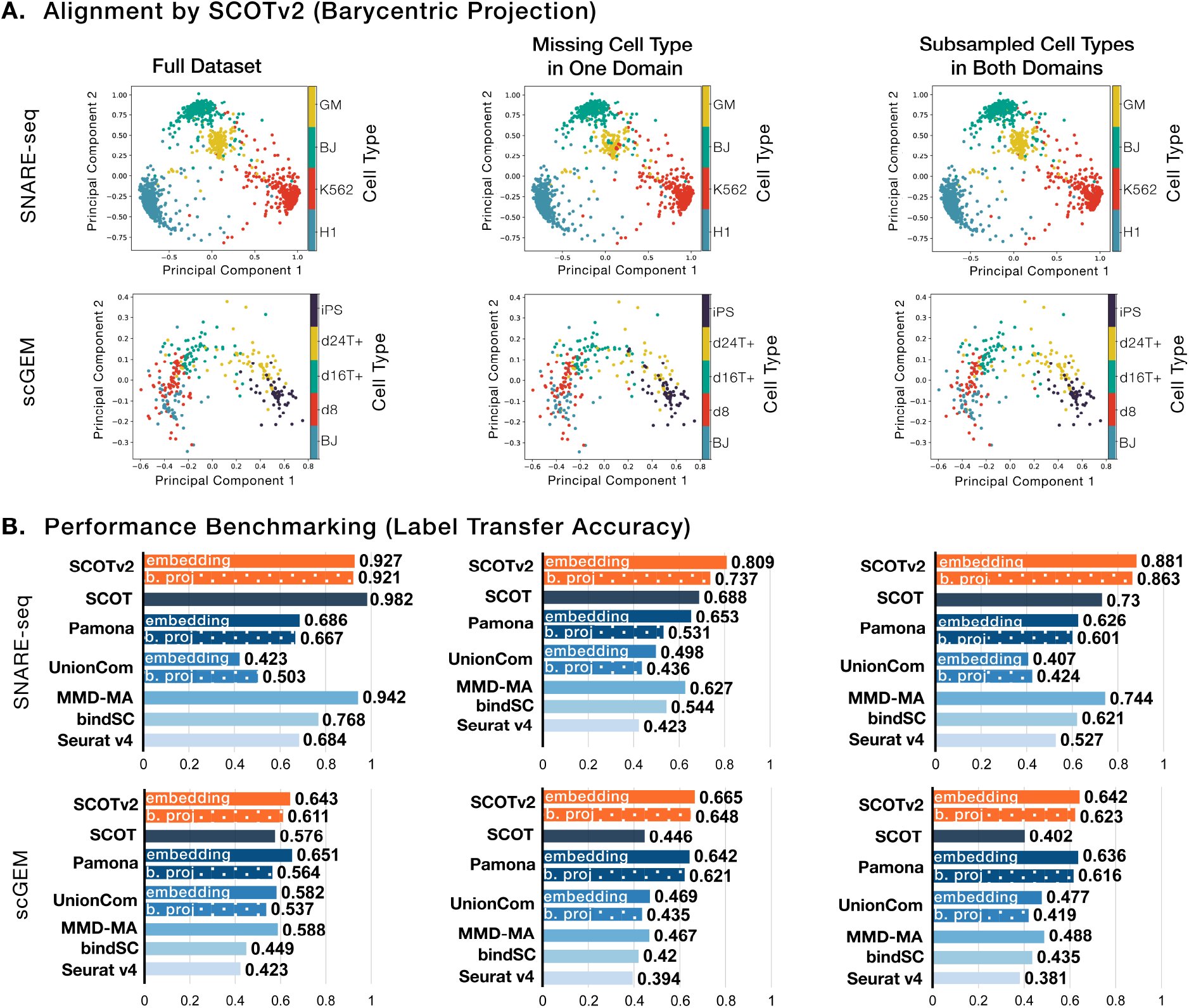
Alignment results for simulations and balanced co-assay datasets. **A** visualizes the barycentric projection alignment on SNARE-seq and scGEM for the full co-assay datasets, simulations with a missing cell-type in the epigenomic domain, and subsampled cell-types in both domains. **B** compares the alignment performance of SCOTv2 to the benchmarks through LTA. For SCOTvs, Pamona, and UnionCom, we report results on both embedding into a shared space (solid bars) and the barycentric projection (dotted bars).

Qualitatively, we see that SCOTv2 preserves the cell-type annotations after alignment for all three settings. In Figure 2 (B), we report the quantitative performance of SCOTv2 and all the other state-of-the-art baselines using the Label Transfer Accuracy (LTA) scores. MMD-MA, UnionCom, Seurat, and bindSC fail to reliably align datasets with disproportionate cell-type representation across modalities. While Pamona tends to yield high-quality alignments for cases with cell-type disproportion, it fails to perform well on the SNARE-seq balanced dataset as well as its subsampling simulation. We additionally apply Pamona to randomly downsampled co-assays (Figure S2). We show that while Pamona’s partial optimal transport framework handles cell-type disproportion better than the balanced optimal transport formulation (demonstrated by SCOT), SCOTv2 still shows an advantage in all SNARE-seq simulations, as well as the smaller downsampling schemes (∼10%).

Among all methods tested, SCOTv2 consistently gives more high-quality alignments across different scenarios of cell-type representation. It also demonstrates a ∼ 22% average increase in LTA over the previous version of the algorithm (SCOT) when comparing the barycentric projection results and ∼27% for the embedding results. Supplementary Figure S2 presents similar results (SCOTv2 attains an LTA of 0.786 followed by Pamona at 0.62 on SNAREseq and 0.542 followed by Pamona at 0.538 on scGEM) for missing cell-types in the other (gene expression) domain, suggesting that our choice of domain with missing cell-type does not affect the performance comparison results. UnionCom, Pamona, and SCOTv2 allow us to perform both barycentric projections and embed the single-cell domains in a lower-dimensional space. Overall, we observe that embedding yields higher LTA values than barycentric projection. Since the barycentric projection projects one domain onto another, the separation of the domain being projected onto (or anchor domain) limits the clustering separation after alignment. In contrast, the embedding utilizes t-SNE to enhance cell-type separation, allowing for better-separated clusters after alignment.

Next, we report the alignment performance of SCOTv2 on single-cell datasets with disparities in celltype representation due to sampling during experiments. We include scNMT, a co-assay with varying levels of cells across domains due to quality control procedures, along with sciOmics and MEC for this experiment. Note that scNMT and sciOmics have three different modalities, and hence, we can only report the baselines for methods that can align datasets with *M* > 2. Figure 3(A) presents the qualitative alignment results for SCOTv2 with PCA. SCOTv2 performs well on all three datasets, including the ones with three modalities. The LTA scores in Figure 3(B) demonstrate that SCOTv2 consistently yields the best alignments on the three real-world datasets. These results highlight its ability to reliably integrate separately sampled with disproportionate cell-type representation and multiple (*M* > 2) modalities simultaneously.

**Figure 3:**
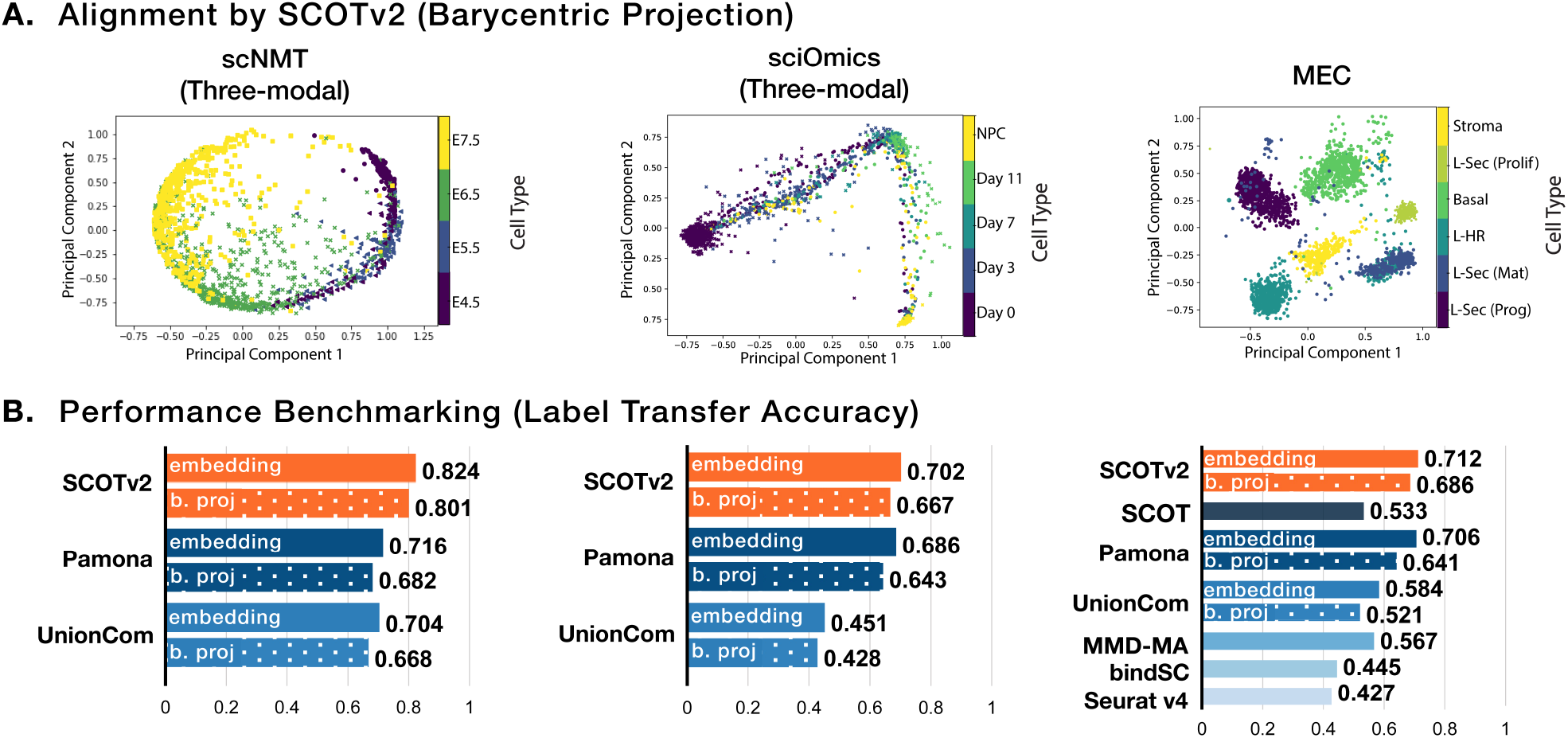
Alignment results for multi-modal (*M* > 2) and separately sequenced datasets. **A** visualizes the alignment of scNMT-seq, sciOmics, and MEC. All datasets have unequal sample sizes and cell-type proportions across domains. **B** benchmarks alignment performance through LTA. As in Figure 2, we report results both by embedding (solid bars) and barycentric projection (dotted bars) for the methods that allow for both. For scNMT-seq and sciOmics, which are three-modal datasets, we only demonstrate results for SCOTv2, Pamona, and UnionCom, which can handle more than two modalities.

### 4.2 Hyperparameter self-tuning aligns well without depending on orthogonal correspondence information

The benchmarking results above present the alignment performance of each algorithm at its best hyperparameter setting; however, users may not have 1—1 correspondences to validate alignments, for the purpose of hyperparameter selection, in real-world applications. While users may have access to cell-type labels, inferring cell-types is highly difficult in specific modalities of single-cell sequencing, such as 3D chromatin conformation. Additionally, different sequencing modalities might disagree on cell-type clustering (as is often the case with scRNA-seq and scATAC-seq datasets). In these situations, users might not have sufficient validation data for tuning hyperparameters.

We design a heuristic process (described in Section 2.4), as done previously for SCOT, that allows SCOTv2 to select hyperparameters in a completely unsupervised manner. Other alignment methods do not provide an unsupervised hyperparameter tuning procedure. Therefore, without validation data, a user would have to use the default parameters. In Table 1, we compare alignment performance for our heuristic against the default parameters of other methods. While our heuristic does not always yield the optimal hyperparameter combination, it does give more favorable results over the default settings of the other methods. Thus, we recommend using it in cases that lack orthogonal information for hyperparameter tuning.

**Table 1:** Alignment performance benchmarking in the fully unsupervised setting. We run SCOTv2 and SCOT using their heuristics to approximately self-tune hyperparameters. We use default parameters for other methods due to a lack of similar procedures for unsupervised self-tuning.

|  | SNARE<br>(full<br>dataset) | SNARE<br>(missing<br>cell-type) | SNARE<br>(subsam.<br>dataset) | scGEM<br>(full<br>dataset) | scGEM<br>(missing<br>cell-type) | scGEM<br>(subsam.<br>dataset) | scNMT | sciOmics | MEC |
| --- | --- | --- | --- | --- | --- | --- | --- | --- | --- |
| **SCOTv2** | 0.826 | **0.653** | **0.751** | **0.509** | **0.521** | **0.415** | **0.727** | **0.537** | **0.584** |
| **SCOT** | **0.852** | 0.572 | 0.588 | 0.423 | 0.323 | 0.314 | N/A | N/A | 0.466 |
| **Pamona** | 0.554 | 0.423 | 0.419 | 0.385 | 0.414 | 0.308 | 0.588 | 0.329 | 0.417 |
| **MMD-MA** | 0.523 | 0.407 | 0.431 | 0.360 | 0.296 | 0.287 | N/A | N/A | 0.233 |
| **UnionCom** | 0.411 | 0.406 | 0.422 | 0.332 | 0.315 | 0.276 | 0.474 | 0.306 | 0.349 |
| **bindSC** | 0.713 | 0.584 | 0.475 | 0.387 | 0.254 | 0.262 | N/A | N/A | 0.412 |
| **Seurat** | 0.503 | 0.477 | 0.428 | 0.408 | 0.377 | 0.329 | N/A | N/A | 0.387 |

### 4.3 SCOTv2 scales well with increasing number of samples

We compare the runtime of SCOTv2 with the top performing methods: Pamona, MMD-MA, UnionCom, and the previous version of SCOT by subsampling various numbers of cells from the MEC dataset. MMD-MA, UnionCom, and SCOTv2 have GPU versions, while Pamona and SCOT only have CPU versions. We run MMD-MA and UnionCom on a single NVIDIA GTX 1080ti GPU with VRAM of 11GB and Pamona and SCOT on Intel Xeon e5-2670 CPU with 16GB memory. We also run SCOTv2 on the same CPU to give comparable results to Pamona’s runtimes. Figure 4 depicts that SCOT, MMD-MA, Pamona, and SCOTv2 show similar computational scaling.

**Figure 4:**
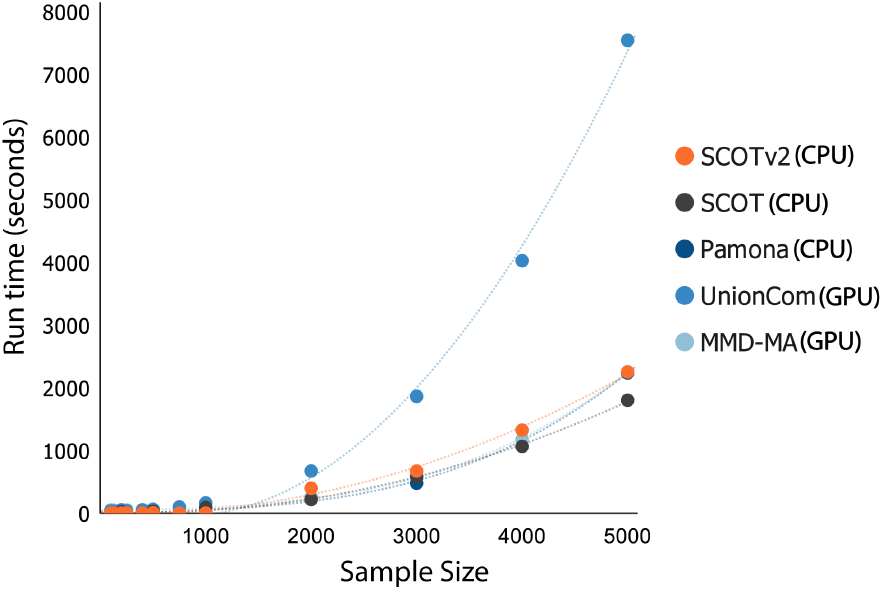
Runtimes for SCOTv2, SCOT, Pamona, UnionCom, and MMD-MA as the number of samples increases.

## 5 Discussion

We present SCOTv2, an improved unsupervised alignment algorithm for multi-omics single-cell alignment. It extends the alignment capabilities of SCOT to datasets with cell-type representation disproportions across different sequencing measurements. It also performs alignment for single-cell datasets with more than two measurements (*M* > 2). Experiments on real-world subsampled co-assay datasets and separately sampled and sequenced single-cell datasets demonstrate that SCOTv2 reliably yields high-quality alignments for a wide range of cell-type disproportions without compromising its computational scalability. Furthermore, SCOTv2’s flexible marginal constraints enable it to consistently give good alignments results for both balanced and unbalanced single-cell datasets. In addition to effectively handling cell-type imbalances and multi-omics alignment, SCOTv2 can self-tune its hyperparameters making it applicable in complete unsupervised settings. Therefore, SCOTv2 offers a convenient way to align multiple single-cell measurements without requiring any orthogonal correspondence information.

In this second iteration of SCOT, we have utilized the coupling matrix in a new way to find a latent embedding space. While this dimension reduction improves cell-type separation, using the coupling matrix directly may offer even more insights into interactions between the aligned domains. Future work will consider how to use the probabilities in the coupling matrix directly for downstream analysis like improved clustering and pseudo-time inference. Though SCOTv2 has runtimes that scale with other methods, it requires *O*(*n*^2^) memory storage for the distance matrices, which may be an issue for especially large datasets. One way to address this limitation would be to develop a procedure to align a representative subset of each domain that can be extended to the entire dataset. Therefore, we will explore this direction to further improve the scalability of SCOTv2.

## Acknowledgements

We thank Ievgen Redko, Ph.D. for useful discussions of optimal transport theory.

## Funding

Ritambhara Singh’s contribution to the work is supported by NIH award 1R35HG011939-01. Bjorn Sandstede was partially supported by NSF awards 1714429 and 1740741. Rebecca Santorella is supported by the National Science Foundation Graduate Research Fellowship under Grant No. 1644760.

## Supplementary Material

### S1 Embedding Method Details

The full details of t-SNE can be found in [17]. For each domain *m*, we compute *P*^*m*^, an *n*_*m*_ × *n*_*m*_ cell-to-cell transition matrix; each entry 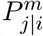 is the conditional probability that a data point 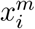 would pick 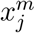 as its neighbor when chosen according a Gaussian distribution centered at 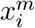:

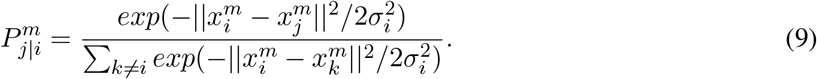

The bandwidth *σ*_*i*_ is chosen according to the density of the data points through a binary search for the value of *σ*_*i*_ that achieves the user-supplied perplexity value.

*P*^*m*^ is computed by averaging 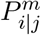 and 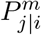 to give more weight to outlier points:

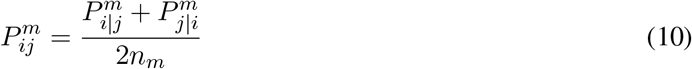

Then, to jointly embed all domains through the anchor domain *X*^1^, the optimization problem is:

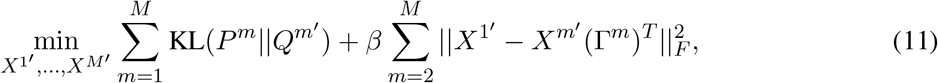

where *X*^*m*′^ is the lower dimensional embedding of *X*^*m*^, *P*^*m*^ is defined as in Equation 9, and Γ^*m*^ is the coupling matrix from solving Equation 6 for *m* = 1, 2, …, *M, X*^*m*′^. The probability matrix *Q*^*m*^ is computed through a Student-t distribution with one degree of freedom:

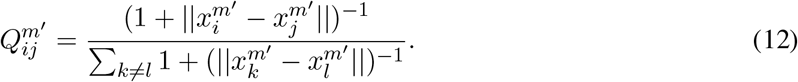

The intuition behind the cost KL(*P*^*m*^||*Q*^*m*′^) is very similar to that of GW; if two points have a high transition probability in the original space, then they should also have a high transition probability in the latent space.

### S2 SCOTv2 Algorithms

#### Algorithm 2: Pseudocode for Unbalanced GW Optimal Transport (UGWOT)

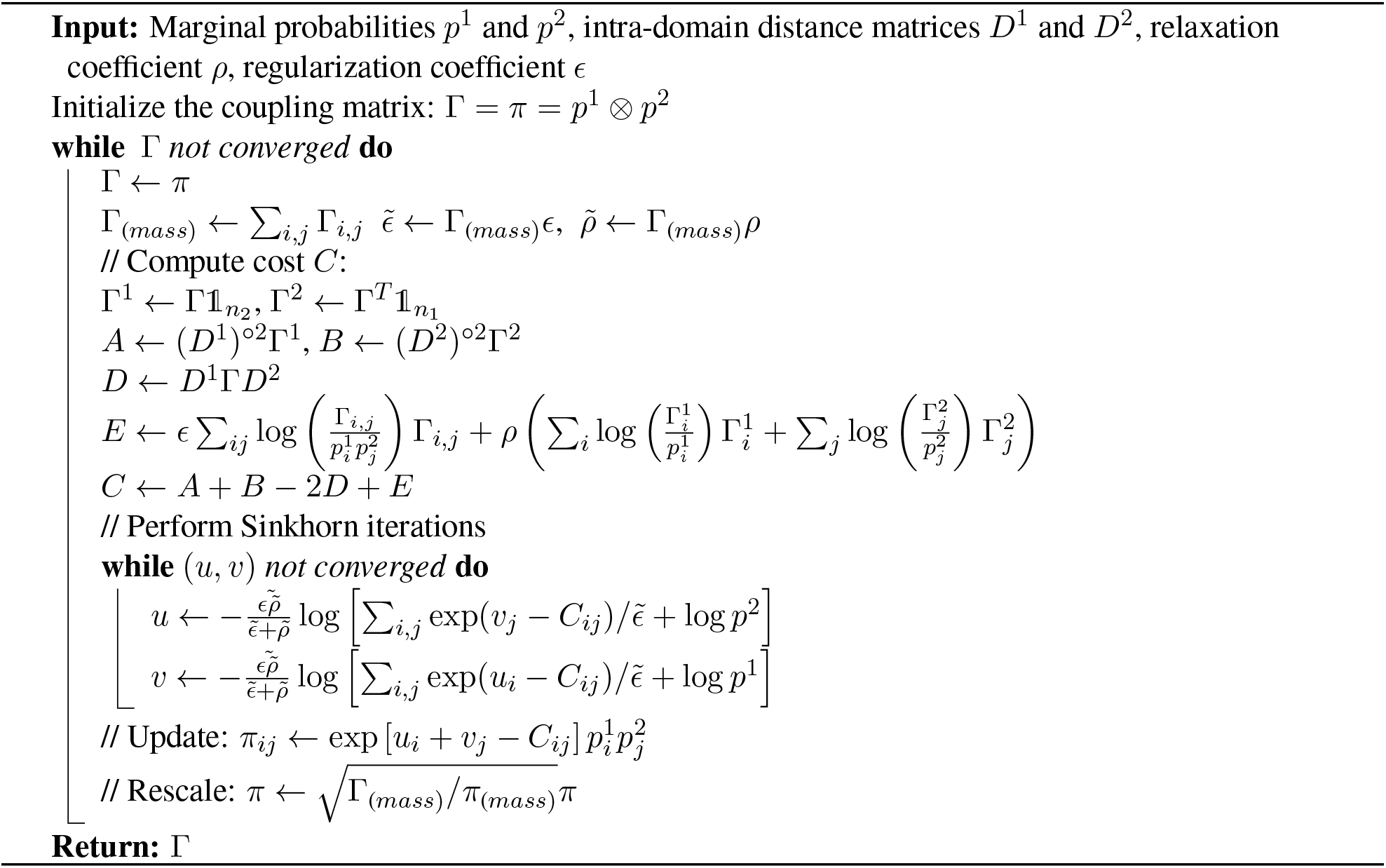

#### Algorithm 3: Pseudocode for SCOTv2 Algorithm

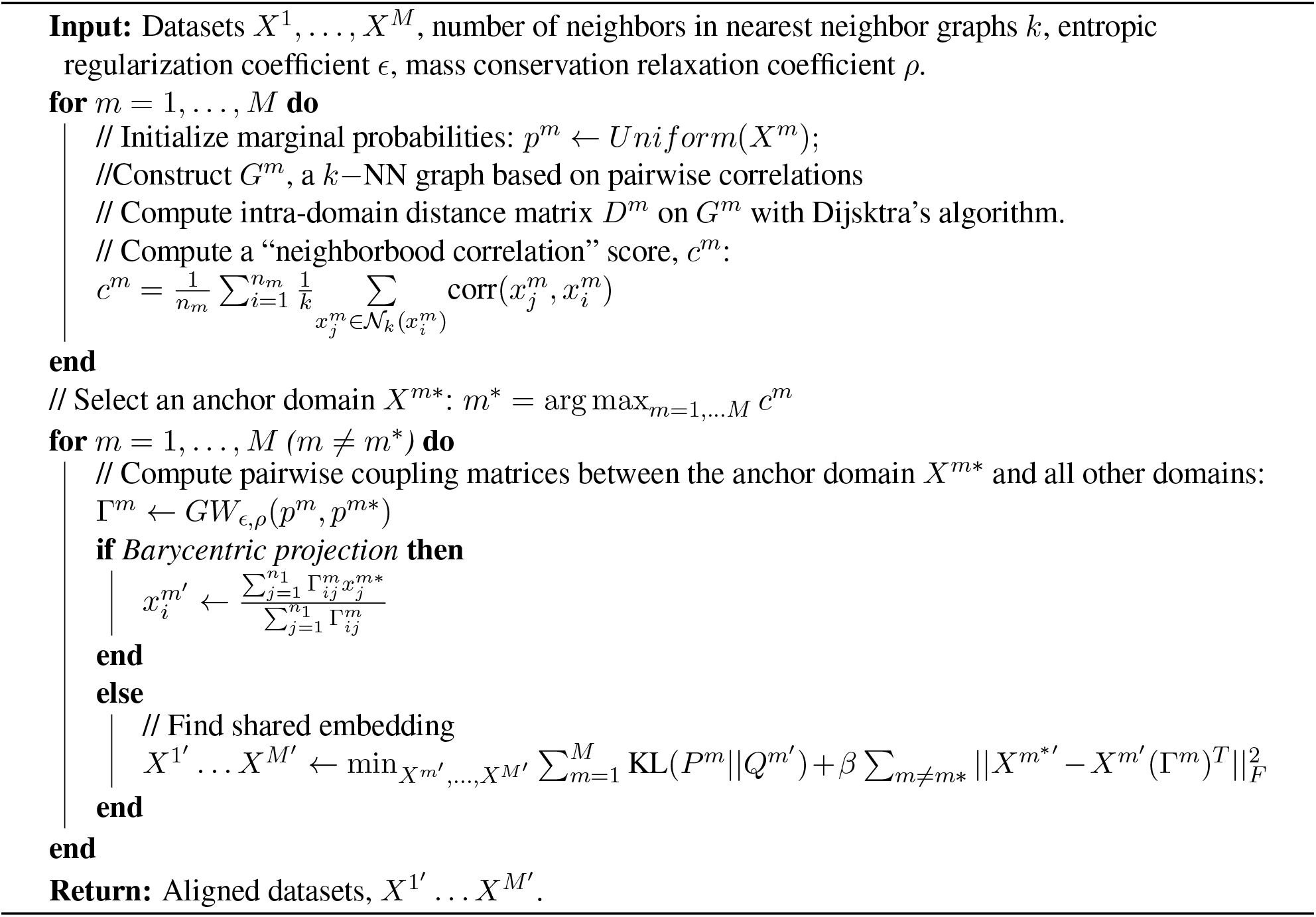

### S3 Cell-type Proportions in Datasets with Intrinsic Cell-Type Representation Disparities across Measurements

**Table S1:** Number of cells in (and percentages of) each cell-type across different modalities in the scNMT-seq co-assayed dataset after quality control procedures and the non-coassay datasets.

|  | **Modality #1**<br>(Gene Expression) | **Modality #2**<br>(Chromatin Accessibility) | **Modality #3**<br>(DNA Methylation or<br>3D chromosomal conform.) |
| --- | --- | --- | --- |
| **scNMT**<br>**dataset** | **(n = 579)**<br>**E4.5:** 76 (12.73%)<br>**E5.5:** 104 (17.42%)<br>**Day6.5:** 146 (24.46%)<br>**E7.5:** 271 (45.39%) | **(n = 647)**<br>**E4.5:** 63 (9.73%)<br>**E5.5:** 89 (13.76%)<br>**E6.5:** 220 (34.00%)<br>**E7.5:** 175 (42.50%) | **(n = 725)**<br>**E4.5:** 65 (8.96%)<br>**E5.5:** 91 (12.55%)<br>**E6.5:** 278 (38.34 %)<br>**E7.5:** 291 (40.14%) |
| **sciOmics**<br>**dataset** | **(n = 1,058)**<br>**Day0:** 489 (46.22%)<br>**Day3:** 127 (12.00%)<br>**Day7:** 78 (7.37%)<br>**Day11:** 145 (13.71%)<br>**NPC:** 219 (20.70%) | **(n = 1,296)**<br>**Day0:** 164 (12.65%)<br>**Day3:** 702 (54.17%)<br>**Day7:** 77 (5.94%)<br>**Day11:** 175 (13.50%)<br>**NPC:** 178 (13.73%) | **(n = 2,154)**<br>**Day0:** 987 (45.82 %)<br>**Day3:** 435 (20.19 %)<br>**Day7:** 243 (11.28 %)<br>**Day11:** 164 (7.61 %)<br>**NPC:** 325 (15.09 %) |
| **MEC**<br>**dataset** | **(n=26,273)**<br>**Basal:** 11,138 (42.39 %)<br>**L-Sec (Prog):** 7,683 (29.24 %)<br>**L-HR:** 3,439 (13.09 %)<br>**L-Sec (Mat):** 2,869 (10.92 %)<br>**L-Sec (Prolif):** 758 (2.89 %)<br>**Stroma:** 386 (1.47 %) | **(n=21,262)**<br>**Basal:** 13,353 (62.80 %)<br>**L-Sec (Prog):** 3,343 (15.72 %)<br>**L-HR:** 2,624 (12.34 %)<br>**L-Sec (Mat):** 1,165 (5.48 %)<br>**L-Sec (Prolif):** 7 (0.033 %)<br>**Stroma:** 770 (3.62 %) | N/A |

### S4 Hyperparameter Tuning Procedure Details

For each alignment method, we define a grid of hyperparameters and choose the best performing combination for each experiment. If methods share similar hyperparameters in their formulation, we keep the range defined for these consistent across all algorithms. Examples for such hyperparameters are dimensionality of the latent space, *p*, for the algorithms that commonly embed datasets; entropic regularization constant, *ϵ*, for methods that employ optimal transport; number of neighbors, *k*, for methods that model single-cell datasets with nearest neighbor graphs. Otherwise, we refer to the publication and the code repository for each method to choose a hyperparameter range.

For Pamona, we tune four hyperparameters: *k* ∈{20, 30, …, 150}, the number of neighbors in the cell neighborhood graphs, *ϵ* ∈ {5*e* − 4, 3*e* − 4, 1*e* − 4, 7*e* − 3, 5*e* − 3, …, 1*e* − 2}, the entropic regularization coefficient for the optimal transport formulation, *λ* ∈ {0.1, 0.5, 1, 5, 10}, the coefficient for the trade-off between aligning corresponding cells and preserving local geometries, and lastly, *p* ∈ {3, 4, 5, 10, 30, 32}, the output dimension for embedding. We choose the ranges for *ϵ* and *k* to be consistent with the corresponding hyperparameters in SCOT and SCOTv2 algorithms and the ranges for the embedding dimensions to be consistent with the recommended values in MMD-MA and UnionCom embeddings.

For UnionCom, we tune the trade-off parameter *β* ∈ {0.1, 1, 5, 10, 15, 20} and the regularization coefficient *ρ* ∈ {0, 0.1, 1, 5, 10, 15, 20} based on the ranges reported by Cao *et al*. in the publication [1]. We additionally tune the maximum neighborhood size permitted in the neighborhood graphs, *k*_*max*_ ∈ {40, 100, 150}, as well as the embedding dimensionality *p* ∈ {3, 4, 5, 10, 30, 32}. The sweep range for hyperparameter *k*_*max*_ is smaller than the other hyperparameters because UnionCom automatically starts from *k* = 2 and goes up to *k*_*max*_ to find the lowest *k* that returns a connected graph to use in the algorithm. Therefore, more refined search is not needed.

For MMD-MA, we choose the weights *λ*_1_ and *λ*_2_ ∈ {1*e* − 2, 5*e* − 3, 1*e* − 3, 5*e* − 4, …, 1*e* − 9}. This range includes the hyperparameter range suggested by Singh *et al* (*λ*_1_, *λ*_2_ ∈ {1*e* − 3, 1*e* − 4, 1*e* − 5, 1*e* − 6, 1*e* − 7}) but extends it further to increase the granularity for the sake of more fair comparison against methods that require a higher number of hyperparameters to test, such as Pamona and UnionCom. Similarly to other methods, we also select the embedding dimensionality from *p* ∈{3, 4, 5, 10, 30, 32}.

For bindSC, we choose the couple coefficient that assigns weight to the initial gene activity matrix *α* ∈ {0, 0.1, 0.2, 0.9} and the couple coefficient that assigns weight factor to multi-objective function *λ* ∈ {0.1, 0.2,, 0.9}. Additionally, we choose the number of canonical vectors for the embdedding space *K* ∈ {3, 4, 5, 10, 30, 32}.

Lastly, for Seurat v4, we tune the number of neighbors to consider when finding anchors, *k* ∈ {5, 10, 15, 20}, dimensions of the final co-embedding space, *p* ∈ {3, 4, 5, 10, 30, 32} and the choice of the reference and anchor domains when finding anchors.

### S5 Visualization of Original Domains Prior to Alignment

**Figure S1:**
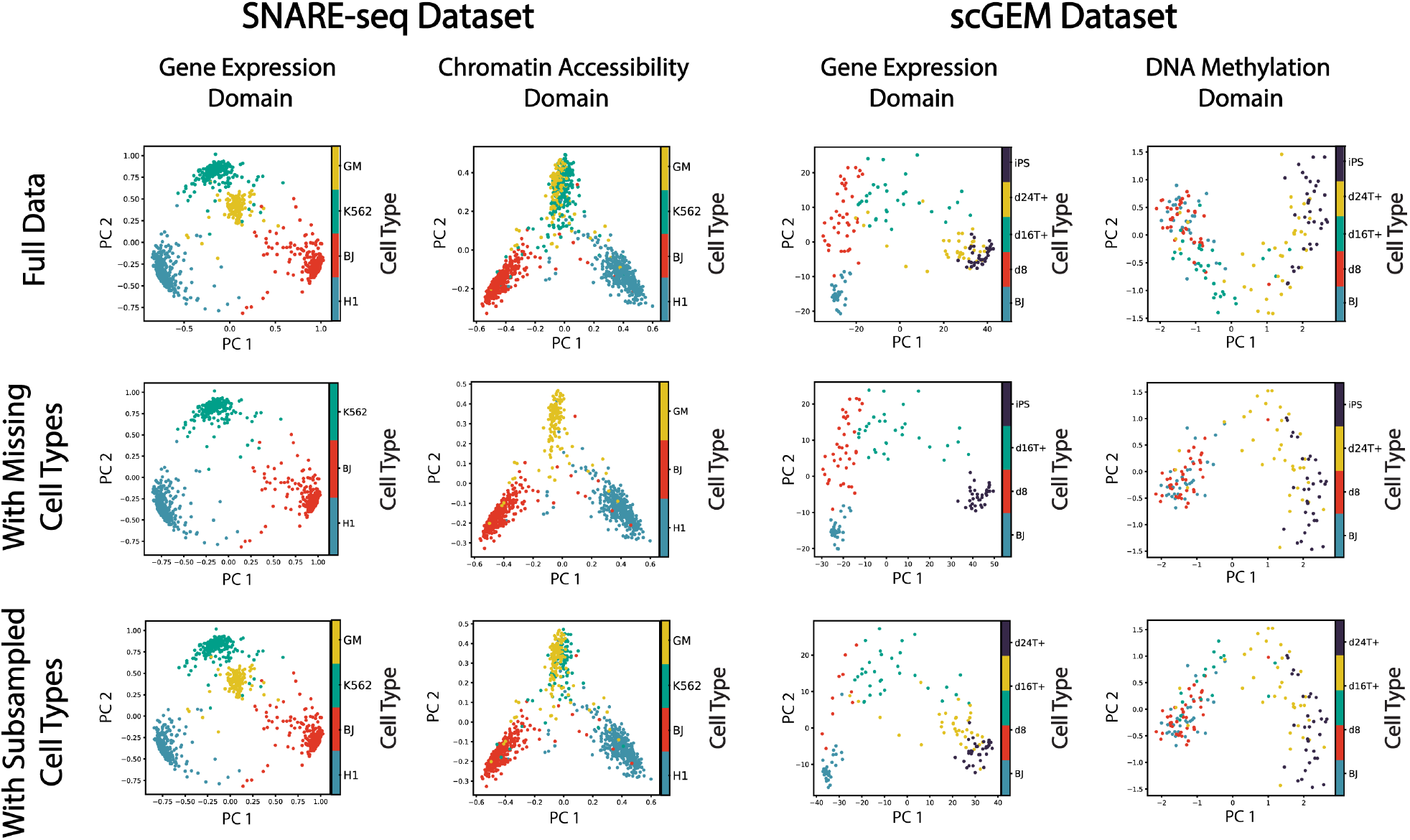
SNAREseq and scGEM datasets prior to alignment for all three simulation scenarios.

### S6 Alignment Results

**Figure S2:**
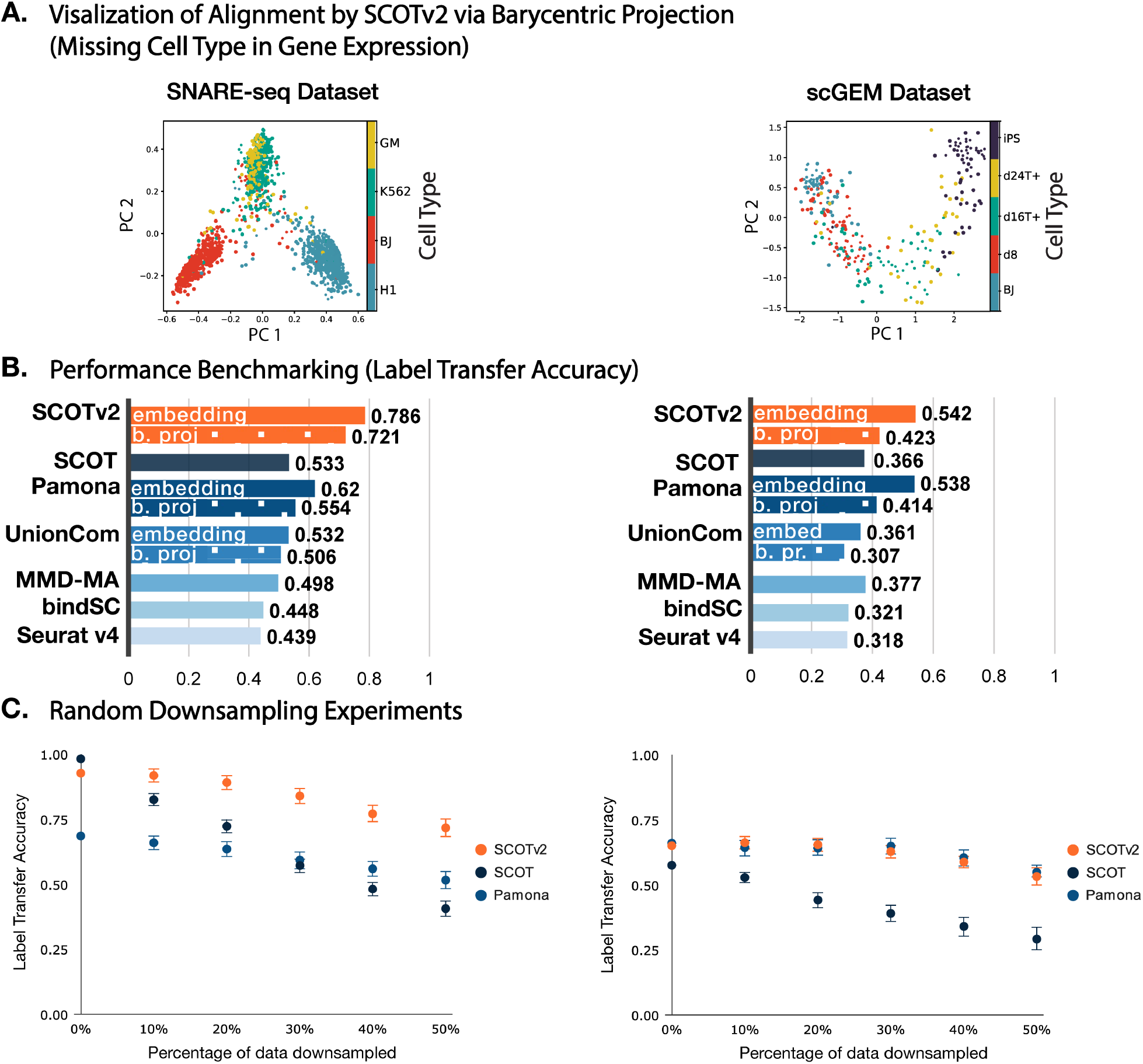
Supplementary alignments results on simulations with co-assay datasets. **Panel A** visualizes the alignment results by SCOTv2, using barycentric projection, on co-assay datasets SNARE-seq and scGEM when a cell-type is missing in the gene expression domain. **Panel B** quantifies the alignment quality in this experiment by using the label transfer accuracy metric and compares to baseline methods. **Panel C** plots the average label transfer accuracy results obtained from SCOTv2, SCOT, and Pamona algorithms when aligning randomly downsampled datasets. These experiments are repeated five times and the standard deviation is shown with error bars.

## Footnotes

1 Preprocessed data for the scGEM dataset accessed here: https://github.com/jw156605/MATCHER

2 Preprocessing scripts for the scNMT-seq data accessed here: https://github.com/PMBio/scNMT-seq/

## Notes

### Competing Interest Statement

The authors have declared no competing interest.

## References

[1] Kai Cao, Xiangqi Bai, Yiguang Hong, and Lin Wan. Unsupervised topological alignment for single-cell multi-omics integration. Bioinformatics, 36(Supplement 1):i48–i56, 2020.

[2] Jie Liu, Yuanhao Huang, Ritambhara Singh, Jean-Philippe Vert, and William Stafford Noble. Jointly embedding multiple single-cell omics measurements. BioRxiv, page 644310, 2019.

[3] Jinzhuang Dou, Shaoheng Liang, Vakul Mohanty, Xuesen Cheng, Sangbae Kim, Jongsu Choi, Yumei Li, Katayoun Rezvani, Rui Chen, and Ken Chen. Unbiased integration of single cell multi-omics data. bioRxiv, 2020.

[4] Tim Stuart, Andrew Butler, Paul Hoffman, Christoph Hafemeister, Efthymia Papalexi, William M. Mauck III, Yuhan Hao, Marlon Stoeckius, Peter Smibert, and Rahul Satija. Comprehensive in-tegration of single-cell data. Cell, 77(7):1888–1902, 2019.

[5] Pinar Demetci, Rebecca Santorella, Bjorn Sandstede, William Stafford Noble, and Ritambhara Singh. Gromov-wasserstein optimal transport to align single-cell multi-omics data. BioRxiv, 2020.

[6] Kai Cao, Yiguang Hong, and Lin Wan. Manifold alignment for heterogeneous single-cell multi-omics data integration using Pamona. Bioinformatics, 08 2021. btab594.

[7] Stephen J Clark, Ricard Argelaguet, Chantriolnt-Andreas Kapourani, Thomas M Stubbs, Heather J Lee, Celia Alda-Catalinas, Felix Krueger, Guido Sanguinetti, Gavin Kelsey, John C Marioni, et al. scnmt-seq enables joint profiling of chromatin accessibility dna methylation and transcription in single cells. Nature communications, 9(1):1–9, 2018.

[8] David Alvarez-Melis and Tommi S Jaakkola. Gromov-wasserstein alignment of word embedding spaces. arXiv preprint 1809.00013, 2018.

[9] Matthias Liero, Alexander Mielke, and Giuseppe Savaré. Optimal entropy-transport problems and a new hellinger–kantorovich distance between positive measures. Inventiones mathematicae, 211(3):969–1117, 2018.

[10] Thibault Séjourné, François-Xavier Vialard, and Gabriel Peyré. The unbalanced gromov wasserstein distance: Conic formulation and relaxation. arXiv, 2021.

[11] Song Chen, Blue B Lake, and Kun Zhang. High-throughput sequencing of transcriptome and chromatin accessibility in the same cell. Nature Biotechnology, 37(12):1452–1457, 2019.

[12] Lih Feng Cheow, Elise T Courtois, Yuliana Tan, Ramya Viswanathan, Qiaorui Xing, Rui Zhen Tan, Daniel S Q Tan, Paul Robson, Loh Yuin-Han, Stephen R Quake, and William F Burkholder. Single-cell multimodal profiling reveals cellular epigenetic heterogeneity. Nature Methods, 13(10):833–836, 2016.

[13] Joshua D Welch, Alexander J Hartemink, and Jan F Prins. Matcher: manifold alignment reveals correspondence between single cell transcriptome and epigenome dynamics. Genome biology, 18(1):138, 2017.

[14] Ricard Argelaguet, Stephen J. Clark, Hisham Mohammed, L. Carine Stapel, Christel Krueger, Chantriolnt-Andreas Kapourani, Ivan Imaz-Rosshandler, Tim Lohoff, Yunlong Xiang, Courtney W. Hanna, Sebastien Smallwood, Ximena Ibarra-Soria, Florian Buettner, Guido Sanguinetti, Wei Xie, Felix Krueger, Berthold Göttgens, Peter J. Rugg-Gunn, Gavin Kelsey, Wendy Dean, Jennifer Nichols, Oliver Stegle, John C. Marioni, and Wolf Reik. Multi-omics profiling of mouse gastrulation at single-cell resolution. Nature, 576(7787):487–491, 2019.

[15] Giancarlo Bonora, Vijay Ramani, Ritambhara Singh, He Fang, Dana L. Jackson, Sanjay Srivatsan, Ruolan Qiu, Choli Lee, Cole Trapnell, Jay Shendure, Zhijun Duan, Xinxian Deng, William S. Noble, and Christine M. Disteche. Single-cell landscape of nuclear configuration and gene expression during stem cell differentiation and x inactivation. Genome Biology, 22(1):279, 2021.

[16] Ritambhara Singh, Pinar Demetci, Giancarlo Bonora, Vijay Ramani, Choli Lee, He Fang, Zhijun Duan, Xinxian Deng, Jay Shendure, Christine Disteche, et al. Unsupervised manifold alignment for single-cell multi-omics data. In Proceedings of the 11th ACM International Conference on Bioinformatics, Computational Biology and Health Informatics, pages 1–10, 2020.

[17] Laurens Van der Maaten and Geoffrey Hinton. Visualizing data using t-sne. Journal of machine learning research, 9(11), 2008.

